# Single particle-resolution fluorescence microscopy of nanoplastics

**DOI:** 10.1101/2020.08.25.267443

**Authors:** Brian Nguyen, Nathalie Tufenkji

## Abstract

Understanding of nanoplastic prevalence and toxicology is limited by imaging challenges resulting from their small size. Fluorescence microscopy is widely applied to track and identify microplastics in laboratory studies and environmental samples. However, conventional fluorescence microscopy, due to diffraction, lacks the resolution to precisely localize nanoplastics in tissues, distinguish them from free dye, or quantify them in environmental samples. To address these limitations, we developed techniques to label nanoplastics for imaging with Stimulated Emission Depletion (STED) microscopy to achieve resolution at an order of magnitude superior to conventional fluorescence microscopy. These techniques include (1) passive sorption; (2) swell incorporation; and (3) covalent coupling of STED-compatible fluorescence dyes to nanoplastics. We demonstrate that our labeling techniques, combined with STED microscopy, can be used to resolve nanoplastics of different shapes and compositions as small as 50 nm. The longevity of the dye labeling is demonstrated in different media and conditions of biological and environmental relevance. We also test STED imaging of nanoplastics in exposure experiments with the model worm *C. elegans*. These techniques will allow more precise localization and quantification of nanoplastics in complex matrices.

**Synopsis:** We show that Stimulated Emission Depletion (STED) microscopy can be used to image single nanoplastics of different compositions and shapes. This will allow researchers to study environmentally-relevant nanoplastics and their interactions with organisms in relevant exposure scenarios.

**TOC Graphic:** 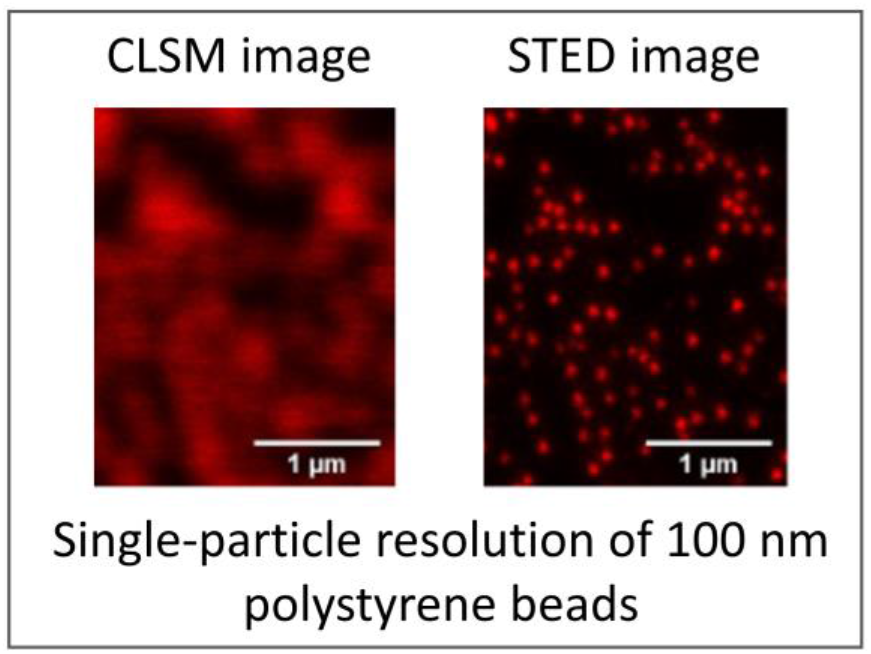

## Introduction

The visibility of plastic pollution and the potential effects on biota have garnered considerable attention. The vast majority of plastics produced ends up in landfills or the environment^1^. Once in the environment, plastic can fragment into smaller pieces known as secondary plastics, via mechanical, thermal, or other mechanisms of degradation ^2–7^. As a result, the smallest size fractions of plastic pollution often dominate particle counts in the environment ^8,9^. Despite their potential prevalence, little is known about the toxicological effects and mechanisms of the smallest particles (i.e., nanoplastics) in part due to the difficulty of visualizing them in wet matrices including biological tissues.

Fluorescence microscopy allows sensitive and quantitative imaging with specific labeling of particles and biological structures. Consequently, fluorescence imaging has been widely applied to localize labeled microplastic particles used in exposure studies to track uptake and potential translocalization in live organisms ^10–12^. Typically, microplastics are large enough to be individually resolved with light microscopy ^10–12^, allowing localization of fluorescently-labeled microplastics even if there is significant background signal from autofluorescence or dye leaching^13,14^. However, because of the optical ~200 nm diffraction limit on resolution, nanoplastics are typically too small to be individually resolved with fluorescence microscopy techniques currently applied in plastics research, including Laser-Scanning Confocal Microscopy^13,15^ and widefield epifluorescence microscopy ^16–18^. Consequently, with conventional light microscopy, nanoplastics are typically only visualized as diffraction limited spots and cannot be precisely localized in tissues nor distinguished from free dye or autofluorescence^13^. While electron microscopy resolution is well beyond the diffraction limit for light microscopy, electron microscopy lacks the labelling flexibility, matrix flexibility, and the ease of sample preparation of fluorescence microscopy.

Nanoplastic exposure experiments disproportionately employ spherical polystyrene nanoparticles, in part due to the limited commercial availability of other types of labeled nanoplastics ^19,20^. These polystyrene spheres are not representative of the diversity of environmentally relevant plastic contamination ^19,20^. Techniques have been developed to label microplastics of arbitrary shapes and compositions ^10^ and Stimulated Emission Depletion (STED) microscopy has been used to image nano-sized polystyrene latex with pre-loaded proprietary fluorophores ^21^. However, methods to fluorescently label nanoplastics of different shapes and polymer types are limited. Consequently, a method to label nanoplastic particles of various shapes and composition would allow more environmentally relevant study of nanoplastics transport, uptake and translocation.

In this work, we developed techniques to label nanoplastics of various shapes, sizes, and polymer types with STED-compatible dyes to image at diffraction-unlimited resolution. We show that nanoplastics can be labeled either by (1) passive sorption; (2) swell incorporation; or (3) covalent coupling of STED-compatible fluorescence dyes. We display STED-compatible labeling and fluorescent imaging of multiple types of nanoplastics, including secondary nanoplastics, of different shapes and polymer types as well as nano-scale resolution imaging of nanoplastics in *Caenorhabditis elegans*, a model nematode worm. We expect that these techniques will extend the application of super-resolution fluorescence microscopy techniques to the localization and identification of nanoplastic translocation at the nano-scale in cells, tissues, and smaller organisms in exposure experiments.

## Materials and Methods

### Nanoplastic sources

Plain polystyrene spheres with a nominal diameter of 100 nm (actual diameter averaged 88 nm according to manufacturer specifications) and 50 nm (actual diameter averaged 50 nm according to manufacturer Dynamic Light Scattering specifications) were purchased from Polysciences Inc. Amine-functionalized 100 nm (actual diameter averaged 113 nm according to manufacturer Dynamic Light Scattering specifications) polystyrene spheres was purchased from Polysciences Inc. Plain 50 nm (actual diameter averaged 50 nm according to manufacturer specifications) Poly(methyl methacrylate) spheres were purchased from Phosphorex Inc. These commercially available spherical particles are typical of those used in most current nanoplastic studies^13,15^. A dispersion of polytetrafluoroethylene (PTFE) nanoparticles (Teflon™ 30B) was purchased from Polysciences Inc. This dispersion has typical average particle diameters around 200 nm but are not monodisperse and contain a variety of particle sizes including much smaller particles.

We also obtained nano-sized debris from plastic labware and consumer items using a variety of weathering methods. Debris was obtained from a polystyrene Petri dish by mechanical abrasion with a 100 nm diamond lapping film (3M) and from an expanded polystyrene plate by submerging it in approximately 1 L of DI water at approximately 90 °C for 7 days and sampling the water by pipetting into a glass vial. These secondary nanoplastic particles represent nanoplastics that can be released from common single-use plastic consumer goods^34–36^.

### Labeling nanoplastics

#### Passive Staining of Nanoplastics

We labeled nanoplastic via passive sorption with Atto 647N (Sigma 04507) (ex. 646 nm / em. 664 nm). We chose to use Atto 647N because of its good STED performance with a 775 nm depletion laser. To passively label nanoplastic, we suspended 0.125% w/v nanoplastic in a solution of 2 mg/L Atto 647N in DI water for 2 h at room temperature (~ 25 °C). The excess dye was removed from the suspension via dialysis using 12-14 kDa molecular weight cutoff (MWCO) dialysis membranes (Frey Scientific) for 7 days with daily water changes in stirred 2 L glass beakers.

#### Swell Incorporation of Dye to Nanoplastics

We also adapted the microplastic dyeing methods developed previously by Karakolis et al.^10^ to label nanoplastics with iDye (Jacquard Products), an inexpensive commercially available fabric dye. While we expected that iDye would be inferior to Atto 647N in terms of fluorescence imaging, we tested it due to the convenience and low cost of labeling with the iDye Poly Blue^10^. Briefly, 0.125% w/v nanoplastics were incubated in a 0.1 mg/mL solution of iDye Poly Blue in DI water heated to 70 °C for 2 h followed by cooling to room temperature. During the heating, the dye is able to diffuse into the polymer matrix due to thermal swelling. Since iDye Poly is only sparingly soluble, the excess dye was removed via centrifugation (12,000*g* for 45 min), washing the dyed particles twice with 25% isopropanol and three times with DI water. We also tested a similar method using Atto 647N in place of iDye Poly Blue. However, for swell incorporation of Atto 647N, we added 10% tetrahydrofuran to the dye solution to aid in the swelling in addition to heating to 70 °C. We added the tetrahydrofuran since Atto 647N is not packaged with dispersants or other agents which are typically included in commercial fabric dye formulations. We were also able to remove excess dye using dialysis since Atto 647N is more water soluble^37^. Dialysis was conducted using a 12-14 kDa MWCO dialysis membranes (Frey Scientific) for 7 days with daily water changes in stirred 2 L glass beakers^12^.

#### Covalent Coupling of Dye to Nanoplastics

We coupled N-hydroxysuccinimide (NHS)-terminated Atto 647N (Sigma 18373) to amine-functionalized plastic by incubating 0.125% w/v nanoplastic in a solution of 2 mg/L Atto 647N in phosphate buffered saline at pH 7.4. At this pH, the NHS group reacts with the amine groups on the surface of the plastic, covalently binding the dye to the plastic^38^. The excess dye was removed from the suspension via dialysis using a 12-14 kDa MWCO dialysis membranes (Frey Scientific) for 7 days with daily water changes in stirred 2 L glass beakers^12^.

### Fluorescence Imaging

We completed all imaging using an Abberior Expert Line STED microscope using an Olympus Plan-Apo 100x/1.40NA Oil immersion objective. We used a 640 nm laser for excitation and a 775 nm laser for depletion. The microscope was equipped with avalanche photodetectors. We performed both standard laser-scanning confocal and STED imaging on the same microscope. All image analysis was done in ImageJ.

### Oil Longevity Testing

To test the longevity of the dye-labeled nanoplastic in a non-polar solvent, we mounted labeled 100 nm (nominal) polystyrene beads in mineral oil (Sigma M5904) on sealed glass microscope slides stored at room temperature (25 °C) in the dark. Specifically, we spread a 5 μL drop of the nanoplastic suspension on glass slides with a pipette tip, allowed the water to dry and then applied a 20 μL drop of mineral oil onto the slide. We then covered the oil with a #1.5 coverslip and sealed the edges with CoverGrip Coverslip Sealant (Biotium, Inc.). Samples were imaged at each time point using identical imaging settings (10% excitation laser power, 50% depletion laser power, 10 nm pixel size, 10 μs dwell time and 1.0 AU pinhole; see Supplementary Table 1 for imaging settings).

### Aqueous Media Longevity Testing

As water is more volatile than the mineral oil we used, we tested longevity in aqueous solutions, including simulated environmental media, in 1 mL suspensions of labeled nanoplastic in glass vials at room temperature (~ 25 °C) or at elevated temperature (40 °C). Pitcher plant fluid was a gift from Brad’s Greenhouse (Vancouver Island, British Columbia, Canada). Soil water was obtained by mixing 200 mL of DI water with 100 g of dry agricultural soil from McGill University’s MacDonald Campus (Sainte-Anne-de-Bellevue, Quebec, Canada) for 48 h and taking the supernatant after the soil was allowed to settle.

At each time point, we mounted a sample from each suspension onto microscope slides for imaging. Specifically, we spread a 5 μL drop of the nanoplastic suspension on glass slides with a pipette tip, allowed the water to dry and then applied a 20 μL drop of water onto the slide. Samples were imaged at each time point using identical imaging settings (10% excitation laser power, 50% depletion laser power, 10 nm pixel size, 10 μs dwell time and 1.0 AU pinhole for 100 nm particles and 20% excitation laser power, 50% depletion laser power, 5 nm pixel size, 10 μs dwell time and 1.0 AU pinhole for 50 nm particles; see Supplementary Table 1 for imaging settings).

### *C. elegans* culture and exposure

*C. elegans* strain KWN117 worms were obtained from the Caenorhabditis Genetics Center at the University of Minnesota and maintained at 23 °C on Nematode Growth Medium (NGM) plates seeded with *E. coli* OP50 as a food source. KWN117 expresses GFP (488 nm excitation/507 nm emission) in the body wall as well as mCherry (587 nm excitation/610 nm emission) in the apical intestinal membrane^28^.

Prior to exposing the worms to nanoplastic, worms were washed from NGM agar plates with 5 mL of M9 buffer and washed with M9 buffer three times by allowing the worms to settle in 15 mL centrifuge tubes for ~ 10 min and replacing the supernatant with fresh M9 buffer. To expose the worms to nanoplastic, the worms were resuspended in 100 μL of M9 buffer containing 100 ppm nanoplastics and then pipetted onto NGM agar plates. Control worms were prepared similarly but were resuspended in M9 buffer with dialyzed dye solution before pipetting on NGM agar plates. We expect that the dye would be largely removed from the solution with this process. For imaging, the worms were washed off the NGM with M9 buffer and immobilized in low-melting point agarose on microscope slides covered with #1.5 coverslips.

## Results and Discussion

### Nanoplastic labeling and imaging at nanoscale resolution

STED microscopy achieves sub-diffraction resolution by scanning a sample with two lasers simultaneously: an excitation laser with a circular spot and an overlapping depletion laser with a donut-shaped spot^16^. The depletion laser suppresses fluorescence emission from the periphery of the excitation laser spot, effectively decreasing the size of the emission spot which improves resolution beyond the diffraction limit. This results in an order of magnitude improvement in resolution. However, the dye compatibility of STED microscopy is limited due to the need for the dye to efficiently undergo stimulated emission and to have high photostability to limit excessive photobleaching^22,23^. Nevertheless, multiple colours of fluorescent dyes compatible with STED microscopy are available such as Star 440 SXP and SeTau 405, and DY-520XL^24^. Still, the suitability of such dyes to labeling nanoplastics of different shapes and polymer types has not yet been established.

We tested labeling with Atto 647N either via passive sorption, heat/solvent swelling^10^ or covalent coupling (**Fig. 1a-c**). We also tested swell labeling with iDye Poly Blue (“iDye”). The methods to label nanoplastic that we describe here all successfully labeled nanoplastic for STED imaging (**Fig. 1d-g**). **Fig. 2** compares identical fields of view imaged with standard laser-scanning confocal imaging with those obtained using STED imaging. As expected, the resolution of the laser-scanning confocal images was diffraction-limited. Individual particles appear as a diffraction-limited spot significantly larger than their actual size. When particles are close to each other, multiple particles appear as a single body rather than distinct particles. In contrast, with STED imaging using Atto 647N, the size and shape of individual nanoplastic particles as small as 50 nm can be resolved as shown by the images (**Fig. 2**) and corresponding point-spread functions (**Supplementary Fig. 1**). Consequently, STED imaging can detect single nanoplastic particles provided that they are in the field of view of the microscope.

**Figure 1.**
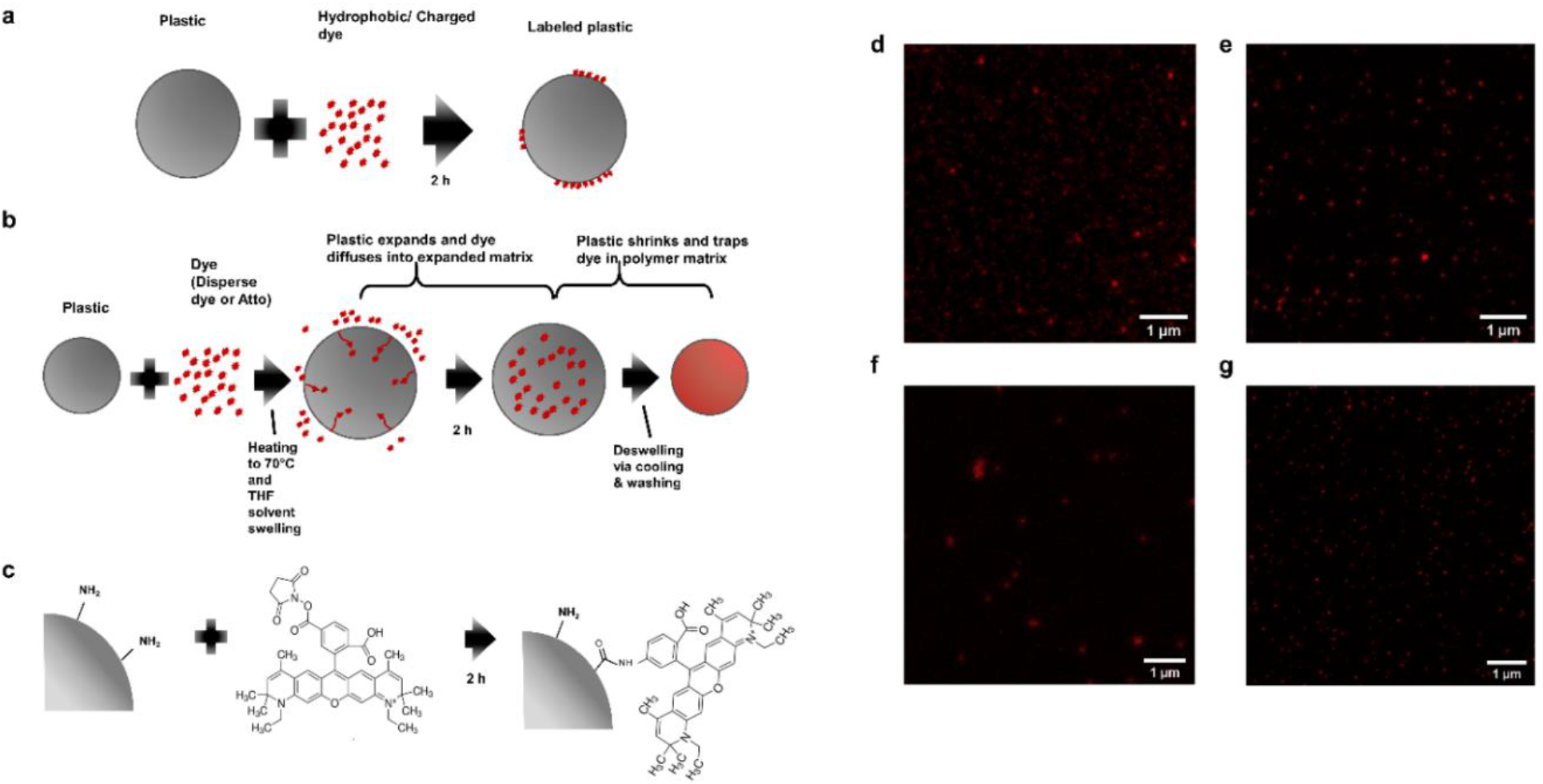
STED-compatible dye labeling techniques (a) Passively-labeling with Atto 647N where the dye solution and the plastic are simply mixed together; (b) Swell-labeling with Atto 647N/iDye Poly Blue where the plastic is heated in the dye solution to swell the polymer matrix and allow the dye to enter the polymer matrix (with Atto 647N, THF, a solvent, is added to aid in swelling), after swelling the plastic is cooled and resuspended in DI water to deswell and remove excess dye; (c) Covalent labeling by coupling NHS-functionalized Atto 647N to amine groups on functionalized plastic; (d) STED image of passively-labeled 100 nm PS beads with Atto 647N; (e) STED image of swell-labeled 100 nm PS beads with Atto 647N; (f) STED image of swell-labeled 100 nm PS beads with iDye Blue; (g) STED image of covalently-labeled 100 nm PS beads with Atto 647N. Dialysis was used to remove excess dye in all cases.

**Figure 2.**
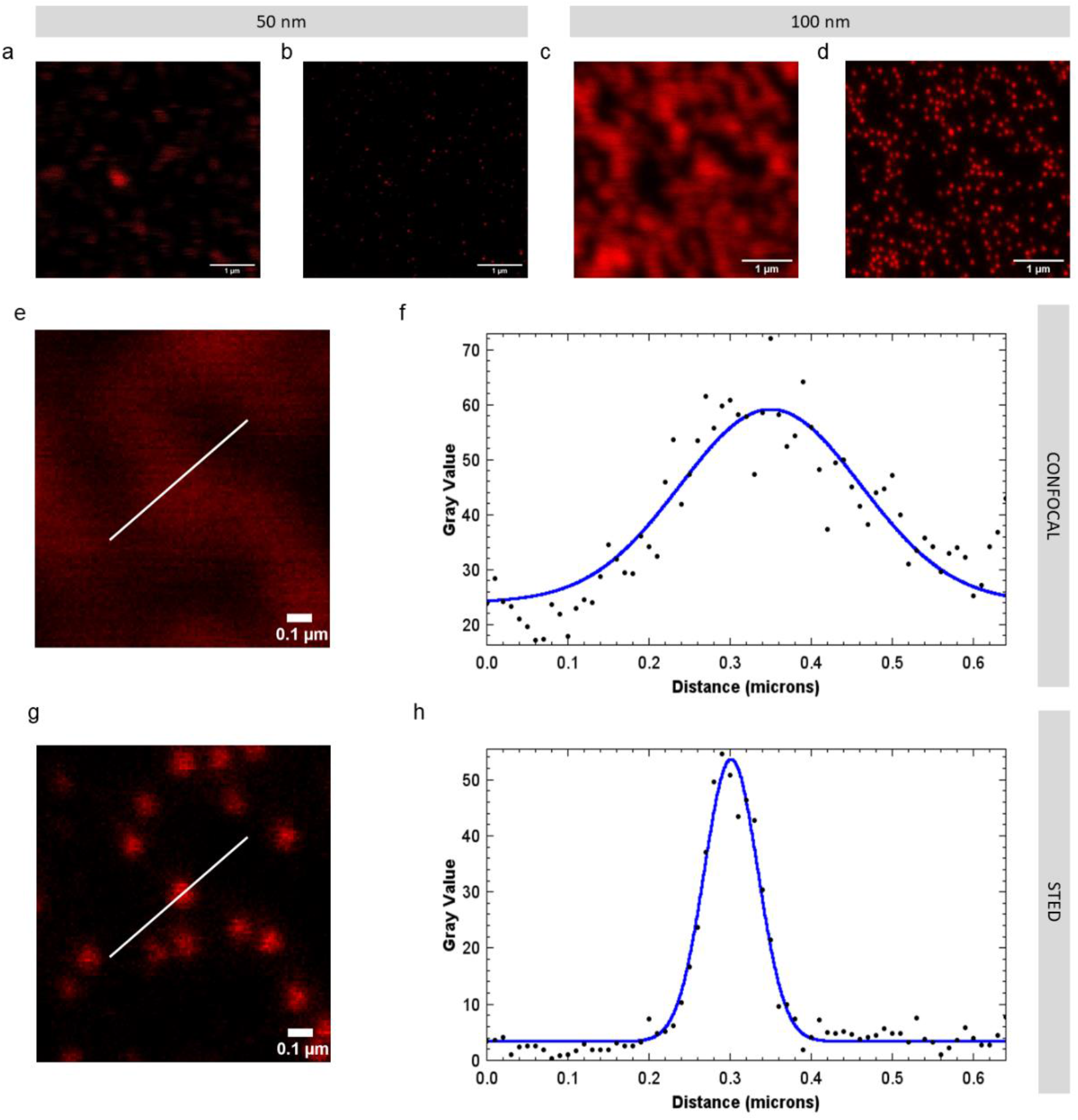
Comparison of Laser-Scanning Confocal images and STED images of identical fields of view where the nanoplastics are passively labeled with Atto 647N. (a) Confocal image of 50 nm polystyrene beads; (b) STED image of 50 nm polystyrene beads; (c) Confocal image of 100 nm polystyrene beads; (d) STED image of 100 nm polystyrene beads; (e) Close up confocal image of 100 nm polystyrene beads; (f) Point spread function and Gaussian fit of 100 nm polystyrene bead with confocal imaging; (g) Close up STED image of 100 nm polystyrene beads; (h) Point spread function and Gaussian fit of 100 nm polystyrene bead with STED imaging.

STED imaging with iDye also showed a resolution improvement over conventional laser-scanning confocal microscopy. The point spread functions of STED images are shown in **Supplementary Fig. 1-4**. However, the resolution with iDye labeling (**Supplementary Fig. 3**) was lower compared to Atto 647N. This result is unsurprising since Atto 647N is well known to perform well with STED microscopy^23^ whereas iDye is a repurposed fabric dye, not a purpose-designed fluorophore. Particles dyed with iDye were also not as bright as those dyed with Atto 647N and thus required 5 times more intense excitation light and twice the pixel dwell time to achieve comparable signal strengths (**Supplementary Table 1**).

### Longevity of Labeled Plastics

We tested the longevity of the labeled nanoplastic particles in both water and oil to simulate the polar and non-polar environments that the particles might experience pre- and post-internalization by organisms. The ability to visualize particles at diffraction-unlimited resolution was maintained in the particles labeled using the techniques we present here for typical exposure timescales. Particularly, all the methods that employ Atto 647N remain visible for at least 49 days (**Fig. 3a-f**). While there was variability day to day in the precise average signal intensity, at all points during this test, the nanoplastic particles were clearly visible with STED imaging throughout the duration of the test. Surprisingly, even plastics labeled passively with Atto 647N were stable in oil (**Fig. 3b**), a non-polar environment. Suspending the plastics in mineral oil did not significantly diminish the fluorescent signal localized to the plastic particle. Moreover, we did not detect a fluorescent signal in the oil phase surrounding plastic particles (directly measured with fluorescence microscopy), indicating that dye transfer to the oil phase is limited.

**Figure 3.**
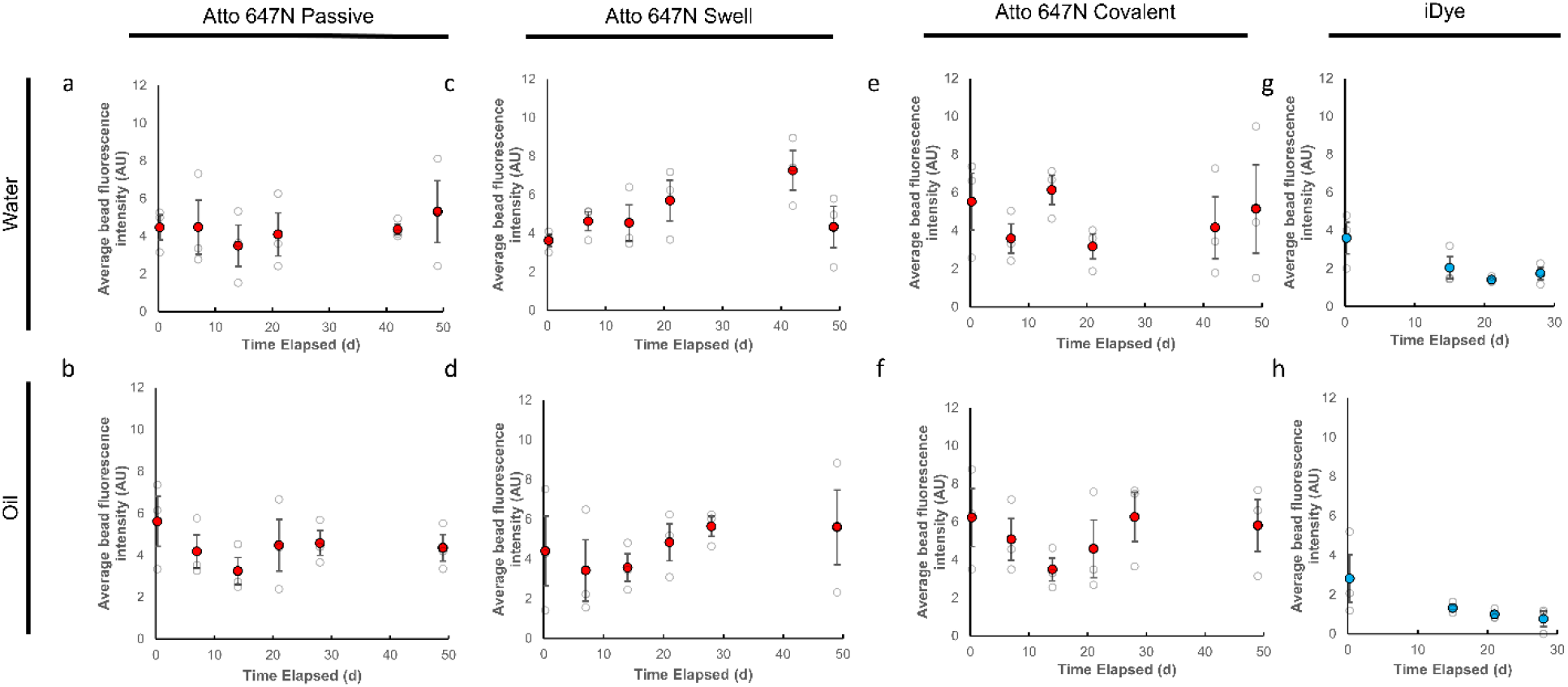
Longevity of different STED-compatible labeling techniques using 100 nm beads. Passive labeling with Atto 647N in (a) DI water and (b) mineral oil; Swell labeling with Atto 647N in (c) DI water and (d) mineral oil; Covalent labeling with NHS-functionalized Atto 647N to amine groups on the particle surface in (e) DI water and (f) mineral oil; Swell labeling with iDye Poly Blue in (g) DI water and (h) mineral oil. Each point represents the average single particle grey value for STED images of triplicate samples acquired with identical settings within treatments. Error bars represent standard error.

The longevity of passively labeled nanoplastics with Atto 647N contrasts with that of Nile Red, a dye more commonly used to stain plastics^11,25^. Previous work^10^ shows that plastics passively stained with Nile Red, do not retain the dye when exposed to mineral oil. This suggests that Atto 647N has a higher affinity to plastic surfaces compared to Nile Red and that the mechanism by which Atto 647N sorbs to plastics is not solely via hydrophobic interaction. Other potential mechanisms of sorption could include interaction between the plastic surface and the charged regions of Atto 647N and/or passive diffusion and subsequent intercalation into the polymer matrix.

To further test the longevity of the labeling in different biologically and environmentally-relevant media and conditions, we also tested the longevity of the different Atto 647N labeling methods on 50 nm polystyrene beads in pitcher plant digestive fluid, soil water, 2.5 pH hydrochloric acid, and at elevated temperature (40 °C) over 21 days (**Fig. 4).** Similar to the tests in water and oil, we found that we were able to visualize the labeled plastics over the period of exposure.

**Figure 4.**
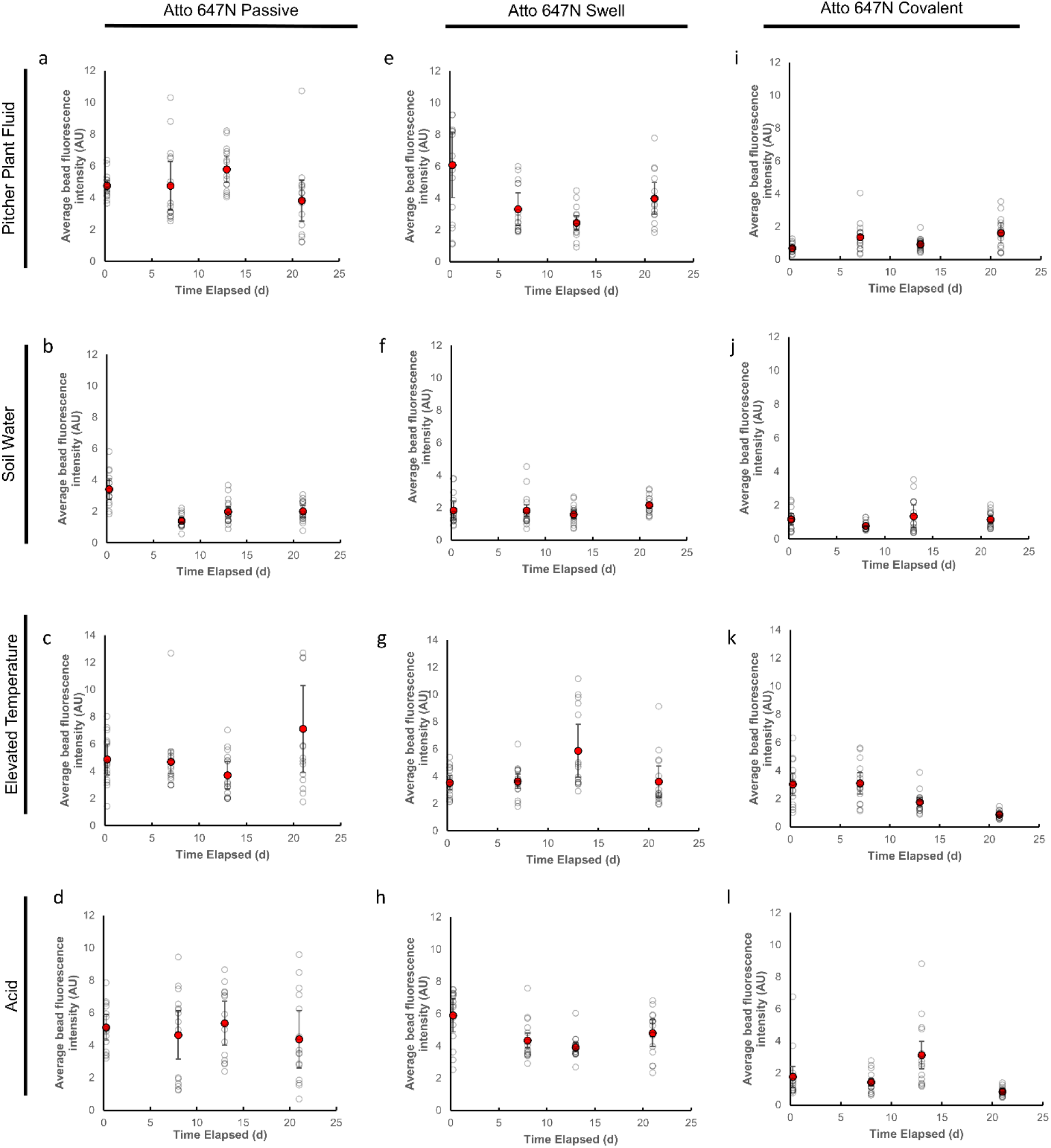
Longevity of different STED-compatible labeling techniques using 50 nm beads. Passive labeling with Atto 647N in (a) pitcher plant fluid; (b) soil water; (c) elevated temperature (40 °C); and (d) hydrochloric acid (pH 2.5). Swell labeling with Atto 647N in (e) pitcher plant fluid; (f) soil water; (g) elevated temperature (40 °C); and (h) hydrochloric acid (pH 2.5). Covalent labeling with Atto 647N in (i) pitcher plant fluid; (j) soil water; (k) elevated temperature (40 °C); and (l) hydrochloric acid (pH 2.5). Each open circle represents the average single particle grey value for STED images of triplicate samples acquired with identical settings within treatments. Error bars represent standard error.

Theoretically, swell and covalent labeling potentially provide greater labeling longevity compared to passive labeling alone. Swell labeling also allows for fluorescent labeling of plastics internally, reducing potential confounds due to surface effects. Nevertheless, in the conditions we tested here, there was little practical difference in the longevity of the particles labeled with different techniques. Moreover, the swell and covalent labeling techniques are comparatively more complicated to carry out. Particularly, with covalent labeling, specific functional groups on the surface of the plastic are required for labeling. Nevertheless, depending on the target application, swell or covalent labeling may be useful when greater resistance to external solvents is desirable.

As with any fluorescent labeling method, the fluorescent signal will be susceptible to photobleaching^10^. Consequently, if experimental exposures require intense light, duration of the exposure while maintaining a fluorescent signal may be more limited. Nevertheless, Atto 647N is generally a photostable dye^26^. Furthermore, despite the resolution advantages that we show, STED microscopy results in a greater degree of photobleaching compared to conventional widefield or confocal fluorescence microscopy since bleaching can also occur via stimulated emission in addition to fluorescence^23^. Consequently, the ability to observe dynamics and image multiple z-slices is limited, in a sample dependent manner, compared to conventional laser-scanning confocal or widefield microscopy. Moreover, to achieve nanoscale resolution, high numerical aperture objectives must be used. Typically, these objectives have limited working distances. Consequently, the maximum imaging depth with STED microscopy while maintaining high resolution is limited and is most suited for imaging cells, tissue sections, small anatomy, or micro-scale organisms.

In terms of performance with STED microscopy, Atto 647N was superior to iDye in brightness, longevity and resolution. Nevertheless, dyeing with iDye may be more practical when cost is a concern. In contrast to Atto 647N, which currently costs $288 CAD per mg, iDye Poly Blue can be purchased for less than $0.50 CAD per g. Nonetheless, typically, a much lower amount of Atto 647N would be required to effectively label nanoplastics compared to iDye Poly Blue.

### Labeling and imaging various types of nanoplastic

While techniques have been developed to label microplastics of arbitrary shapes and compositions^10^, methods to fluorescently label nanoplastics of different shapes and polymer types are limited. Addressing this limitation, our methods are compatible with different shapes and compositions of nanoplastic, beyond commercially available spherical nanoplastics (**Fig. 5**). With STED microscopy, we were able to resolve the shapes of secondary nanoplastics produced by heating expanded polystyrene to 90 °C for 7 days (**Fig. 5b**) as well as polystyrene sanding debris (**Fig. 5d**). As confirmed by SEM (**Supplementary Fig. 9**), the secondary nanoplastics produced by heating are roughly spherical while those produced by sanding are irregular in shape. The images in **Fig. 5a,c** show that such particle visualization was not possible with conventional confocal fluorescence microscopy. We also tested imaging with commercially-available PTFE nanoparticles (**Fig. 5e,f**) as a low surface energy plastic and PMMA nanoparticles (**Fig. 5g,h**) as a relatively high surface energy plastic^27^. The PTFE particles are representative of engineered PTFE nanoparticles used for lubrication and hydrophobic coatings. PMMA (also known as “Plexiglas”) is widely used as a glass substitute for transparent windows.

**Figure 5.**
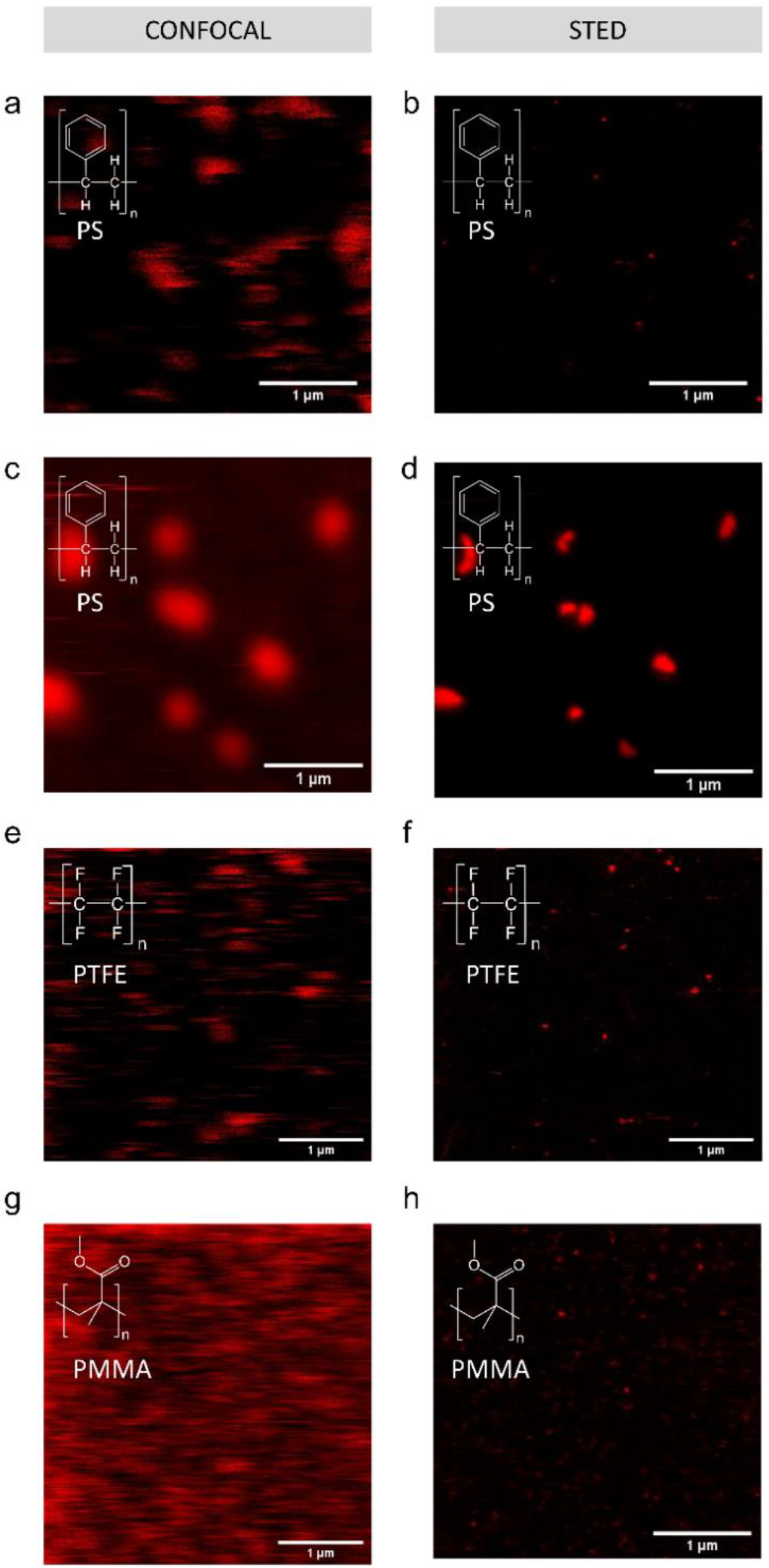
Passive labeling and fluorescent imaging of various nanoplastic types with Atto 647N. Confocal (a) and STED (b) image of debris released from an expanded polystyrene plate exposed to 90 °C in DI water; (c) Confocal and (d) STED image of sanding debris from a polystyrene Petri dish; (e) Confocal and (f) STED image of PTFE particles; (g) Confocal and (h) STED image of PMMA particles. Confocal and STED images are of the same fields of view.

### Application to *C. elegans* exposure

To test the ability of this technique to image nanoplastics in a model organism, as a proof of concept, we exposed *C. elegans* KWN117^28^ to 100 ppm of 50 nm polystyrene beads passively labeled with Atto 647N (**Fig. 6**). We used the 50 nm polystyrene beads due to their availability in sufficient quantities to conduct the exposures as well as a more challenging size to detect. KWN117 is a transgenic strain that expresses Green Fluorescent Protein (GFP) in the body wall and mCherry in apical intestinal membrane cells^28^. As when imaging the nanoplastics alone, STED imaging allows visualization of nanoplastic along sections of the digestive tract including the mouth (**Fig. 6a,b,c**), pharynx (**Fig. 6d,e,f**), and intestine (**Fig. 6g,h,i**). While 100 ppm is higher than typical plastic concentrations for aquatic environments, experiments conducted at higher concentrations are suited to achieving mechanistic insights. Moreover, these concentrations are in line with concentrations in some soil environments ^29,30^.

**Figure 6.**
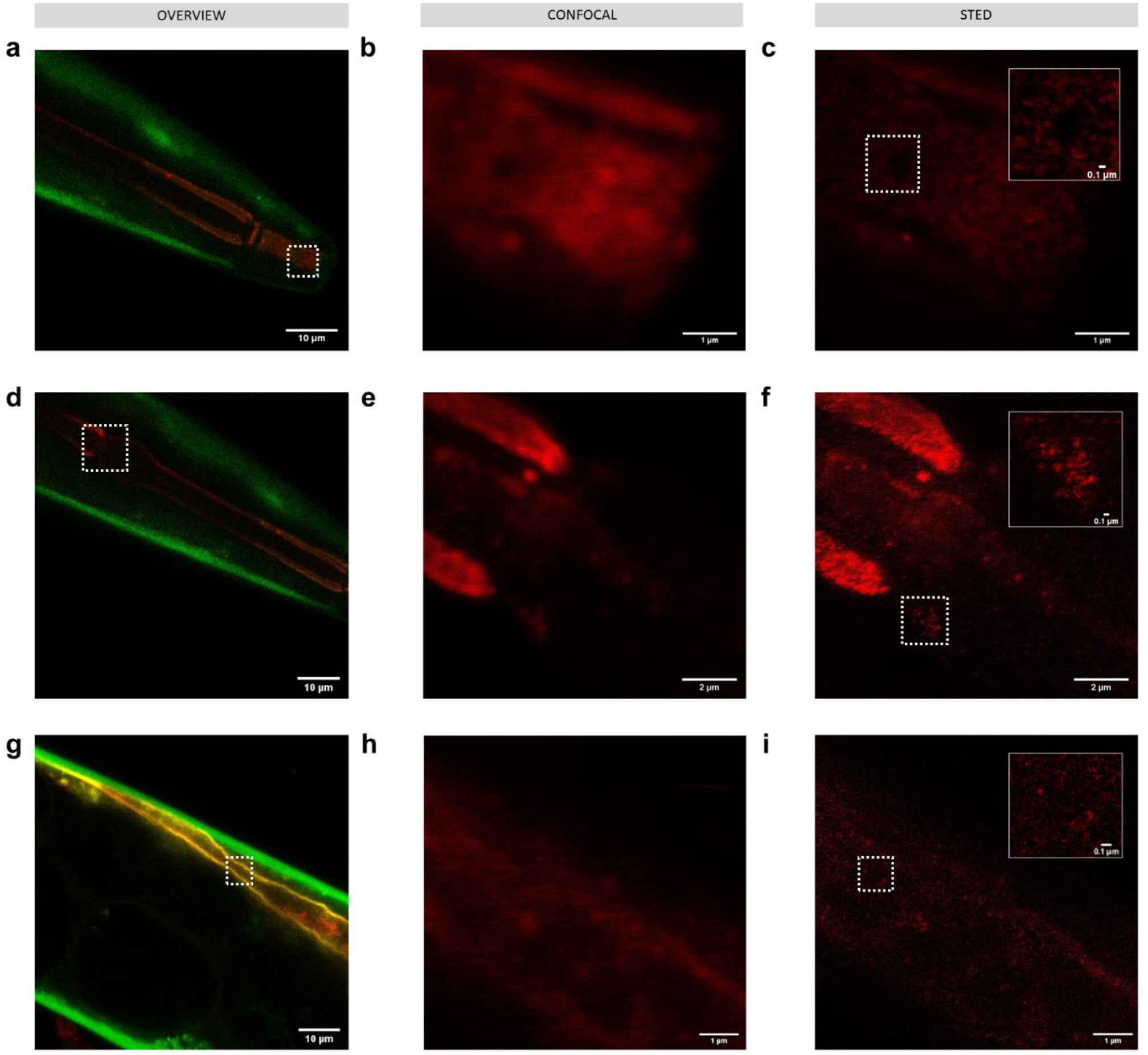
Imaging 50 nm polystyrene nanoplastics passively labeled with Atto 647N (red) in *C. elegans* KWN117 adult expressing GFP (green) in the body wall and mCherry (yellow) in the apical intestinal membrane. Confocal overview, and high resolution of confocal and STED images of scanned area indicated by white boxes for parts of the digestive track in the mouth (a,b,c), pharynx (d,e,f), and intestine (g,h,i). Insets in STED images correspond to the areas indicated by white boxes.

Notably, fluorescent signal in the confocal images, which may initially be interpreted as nanoplastic, is revealed to be background signal from free dye and/or autofluorescence with STED imaging. In control worms (**Supplementary Fig. 8**), no nanoplastic particles were visible with STED imaging, however spots of background and autofluorescence signal were visible that could have been misidentified as nanoplastic with lower-resolution imaging.

### Environmental Implications

We expect that for typical organism exposure experiments, including the imaging in *C. elegans* we show here, STED microscopy combined with passive staining with Atto 647N would generally be a convenient method of observing nanoplastics fluorescently without significantly compromising labeling longevity. Overall, the ability to precisely localize the distribution of nanomaterials, including nanoplastics, in organisms may help facilitate study into the lethal and sub-lethal effects observed in exposure experiments.

Localization can also provide insight into longer term effects of exposure that may not be detected over relatively short experimental timescales. Specifically, translocation of materials to certain parts of organisms can be indicative of long-term effects. Importantly, our results show that STED imaging can detect single nanoplastic particles provided that they are in the field of view of the microscope. Since our methods are compatible with different shapes and compositions of nanoplastic, these techniques will allow researchers to more thoroughly explore the impact of shape, size, and composition of nanoplastics on toxicity and translocation in live organisms. In this work, we demonstrate STED-compatible labeling and fluorescent imaging of different nanoplastics, including secondary nanoplastics, of varying shapes and polymer types. Future work will aim at extending the types of plastics tested, including polyethylene, over a wider range of exposure concentrations. The broad compatibility of STED microscopy to localize fluorescently-labeled nanomaterials at nanoscale resolution can be further extended to better understand the environmental toxicology of nanomaterials beyond nanoplastics. While STED microscopy has been applied to understand the interaction of engineered nanomaterials with cells *in vitro* in the biomedical context^31–33^, our work demonstrates the utility of STED microscopy to study nanomaterial interactions with organisms in environmentally-relevant exposure scenarios and with environmentally-relevant contaminants.

## Acknowledgments

The authors acknowledge Dr. Rachel Genthial, Kamila Mustafina, Dr. Thomas Stroh, Dr. Paul Wiseman, the Integrated Quantitative Biology Initiative, and the Montreal Neurological Institute Microscopy Unit for assistance with STED microscopy. The authors also acknowledge Amin Valiei and David Liu for assistance with SEM. *C. elegans* was provided by the CGC, which is funded by NIH Office of Research Infrastructure Programs (P40 OD010440). B.N. gratefully acknowledges funding from the Natural Sciences and Engineering Research Council Postdoctoral Fellowships program and the Eugenie Ulmer Lamothe fund in the Department of Chemical Engineering at McGill University. N.T. acknowledges funding from the Canada Research Chairs program, the Killam Research Fellowship, the Natural Sciences and Engineering Research Council of Canada, and the Canada Foundation for Innovation. This Project was supported partially by a financial contribution from Fisheries and Oceans Canada.

## ASSOCIATED CONTENT

### Supporting Information Available

Fig. S1-S4 show images and point spread functions of different labeled plastic particles. Fig. S5, S6 show comparison of images of labeled polystyrene beads after 49 days in water and oil. Fig S7 shows an image of the free dye dialyzed control, Fig. S8 shows images of *C. elegans* KWN117 adult exposed to dialyzed dye solution, and Fig. S9 shows SEM images of plastic samples. Table S1 lists the STED imaging settings.

### Author contributions

B.N. developed the methodology and performed the experiments. B.N. and N.T conceived the concepts and contributed to manuscript drafting and editing.

### Data availability

The data that support the findings of this study are available from the corresponding authors upon reasonable request. Figures 3, 4 and Supplementary Figures 1-4 have associated raw data.

