## Supplemental Information for "Single particle-resolution fluorescence microscopy of nanoplastics"

**Supplementary Table 1:** STED imaging settings

| **Sample** | **640 nm excitation laser power** | **775 nm depletion laser power** | **Pixel Size** | **Dwell Time** | **Pinhole** |
| --- | --- | --- | --- | --- | --- |
| **50 nm polystyrene for testing resolution** | 20% | 100% | 10 nm | 10 μs | 1.0 AU |
| **100 nm polystyrene for testing resolution** | 10% | 50% | 10 nm | 10 μs | 1.0 AU |
| **Atto 647N-labeled longevity testing for 100 nm beads (oil and water)** | 10% | 50% | 10 nm | 10 μs | 1.0 AU |
| **Atto 647N-labeled longevity testing for 50 nm beads (elevated temperature, pitcher plant fluid, acid, and soil water)** | 20% | 50% | 5 nm | 10 μs | 1.0 AU |
| **iDye-labeled**  **plastic longevity testing (oil and water)** | 50% | 100% | 10 nm | 20 μs | 1.0 AU |
| **Expanded polystyrene debris** | 40% | 50% | 10 nm | 10 μs | 1.0 AU |
| **Polystyrene sanding debris** | 5% | 35% | 10 nm | 10 μs | 1.0 AU |
| **50 nm PMMA** | 10% | 50% | 10 nm | 10 μs | 1.0 AU |
| **PTFE suspension** | 10% | 50% | 10 nm | 10 μs | 1.0 AU |
| ***C. elegans*** | 35% | 75% | 10 nm | 10 μs | 1.0 AU |

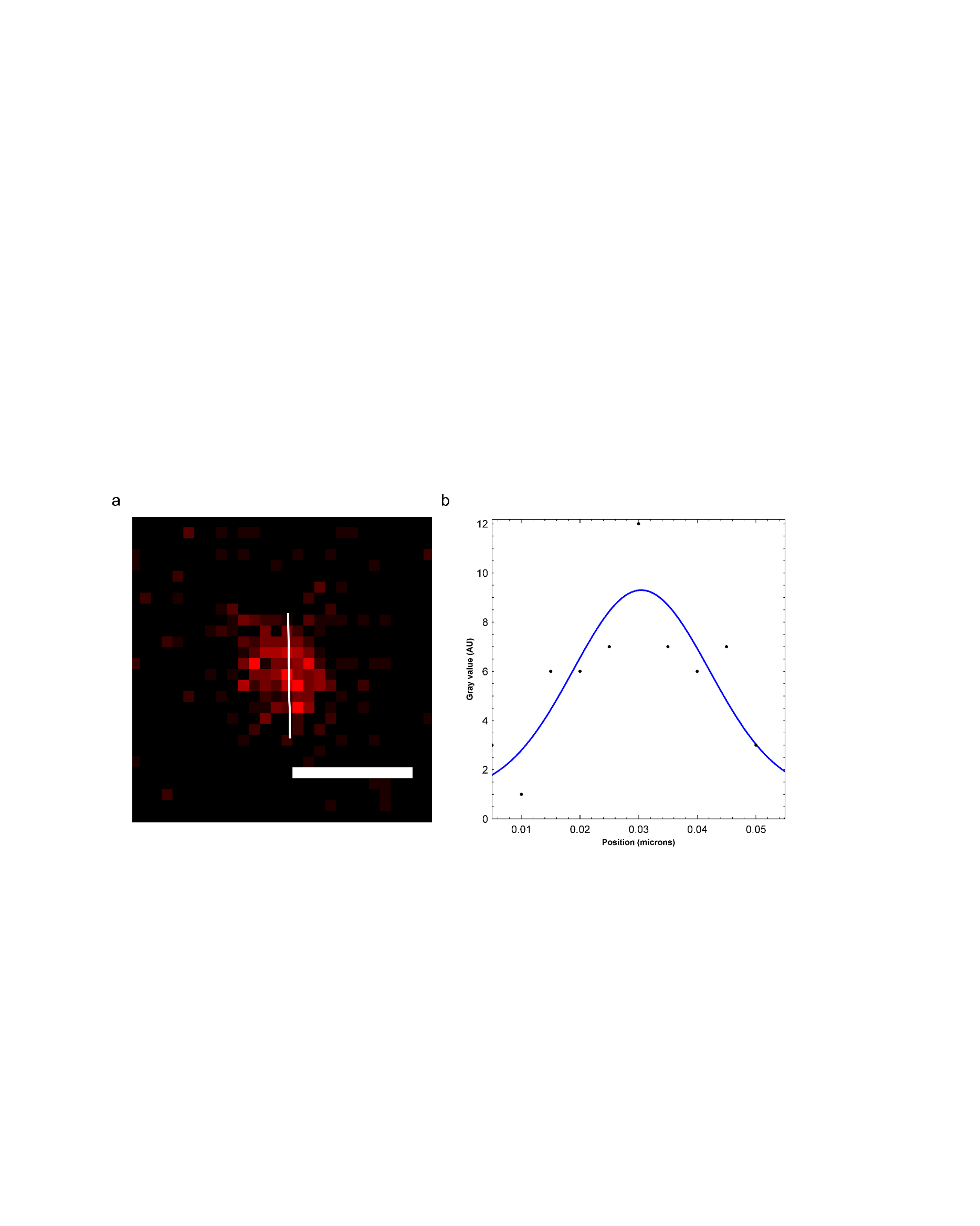

**Supplementary Figure 1:** Image (a) and (b) point spread function across indicated line (Gaussian fit) of a 50 nm polystyrene bead passively labeled with Atto 647N. Scale bar is 50 nm.

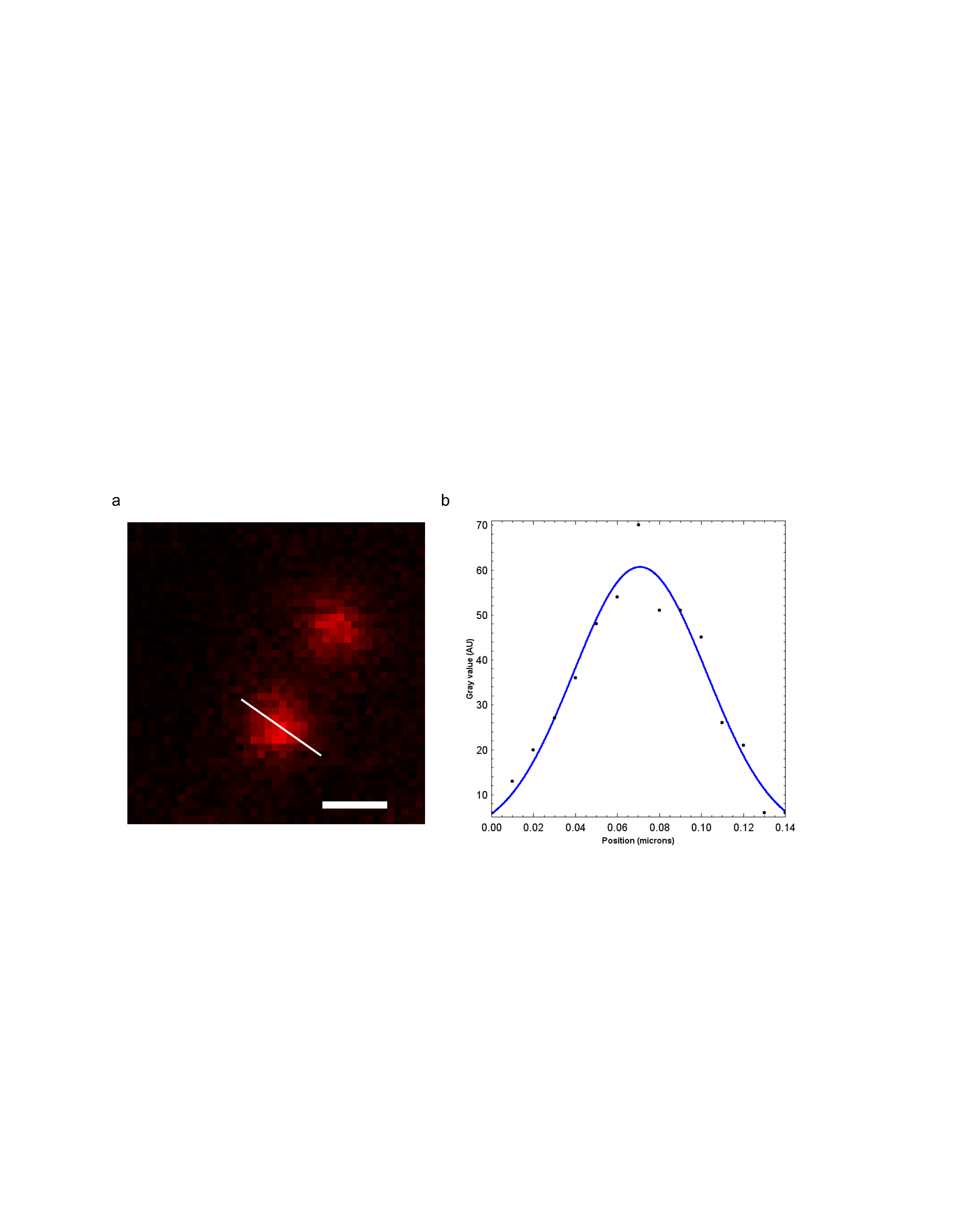

**Supplementary Figure 2:** Image (a) and (b) point spread function across indicated line (Gaussian fit) of a 100 nm polystyrene bead passively labeled with Atto 647N. Scale bar is 100 nm.

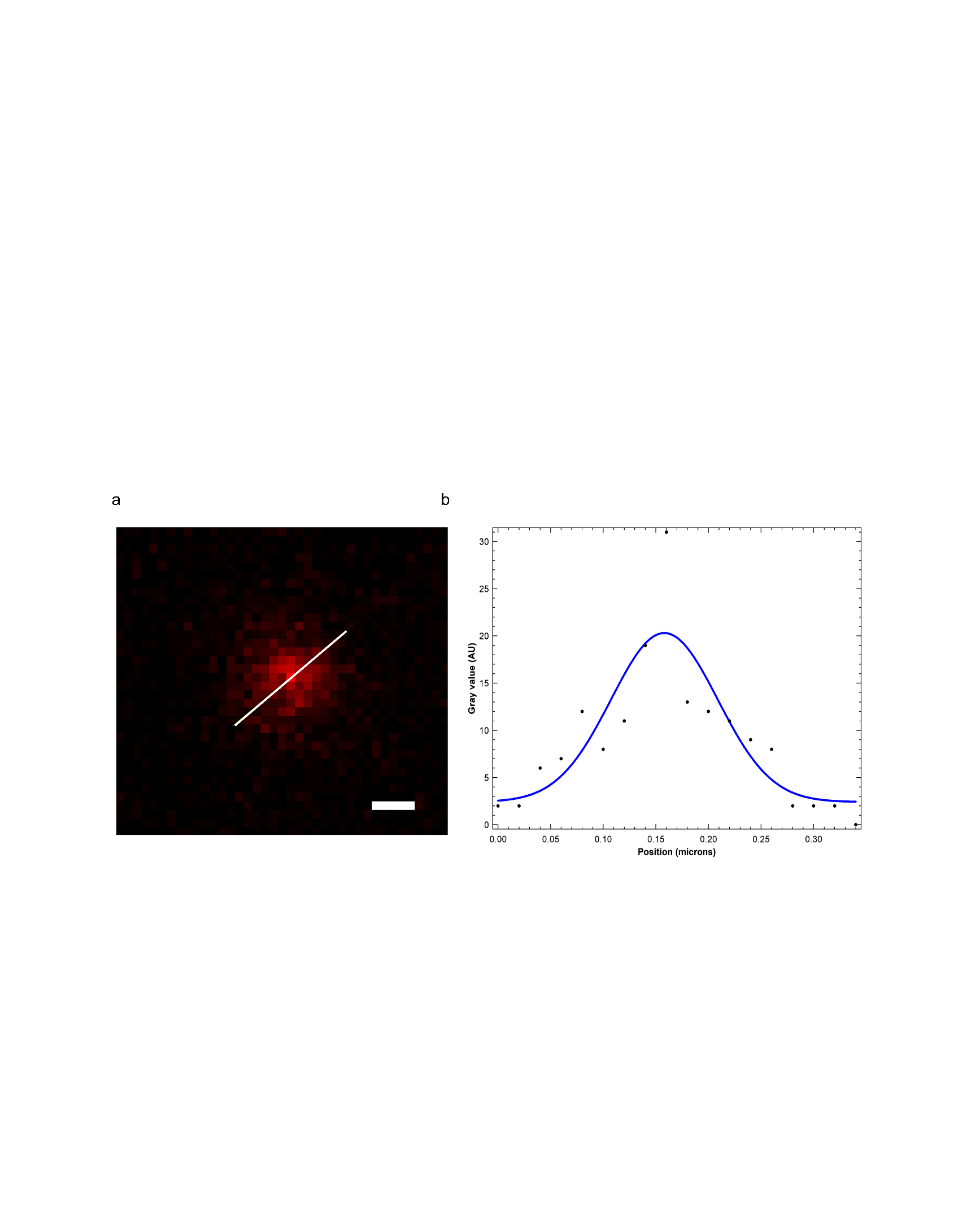

**Supplementary Figure 3:** Image (a) and (b) point spread function across indicated line (Gaussian fit) of a 100 nm polystyrene bead swell labeled with iDye Blue. Scale bar is 100 nm.

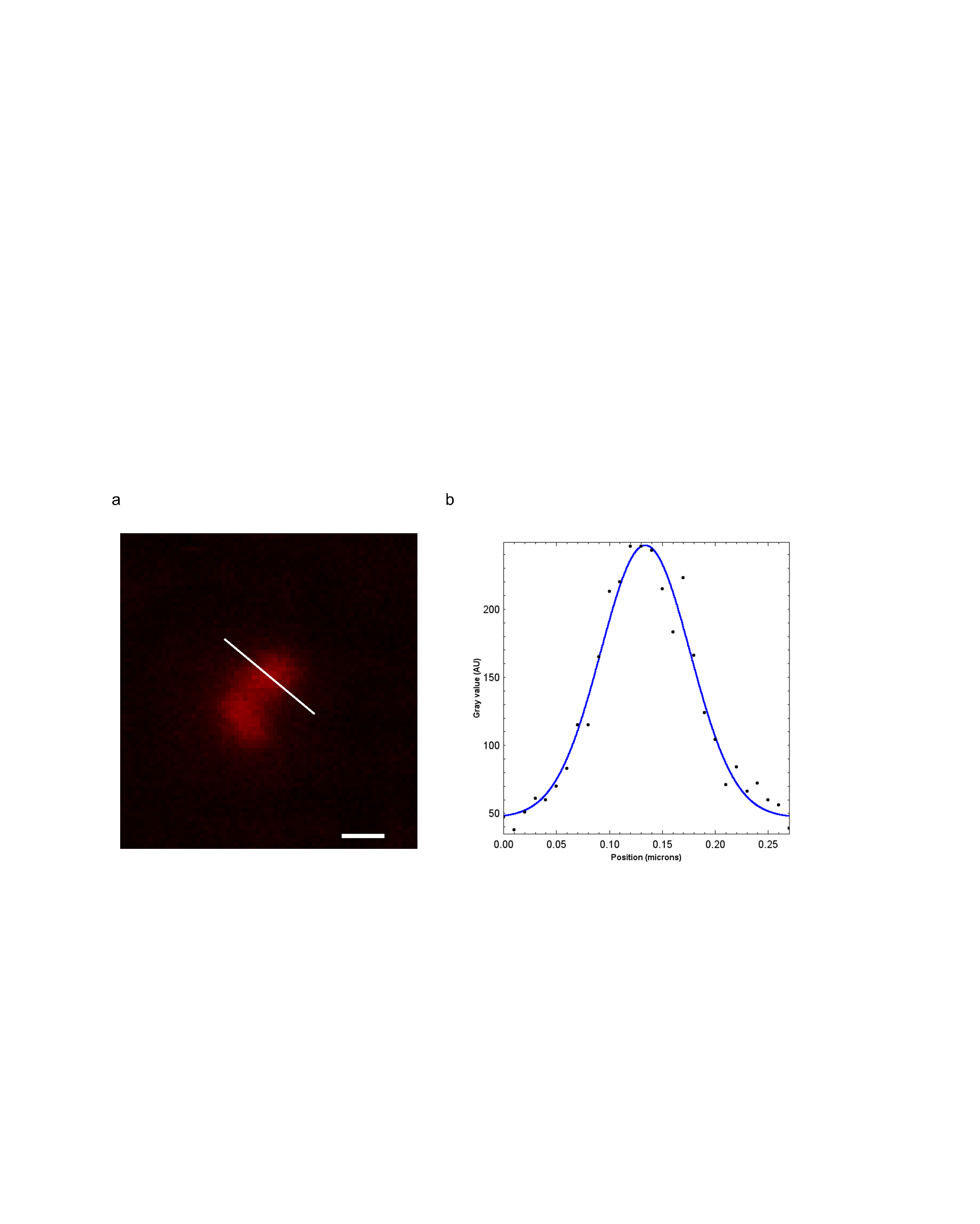

**Supplementary Figure 4:** Image (a) and (b) point spread function across indicated line (Gaussian fit) of a polystyrene sanding debris passively labeled with Atto 647N. Scale bar is 100 nm.

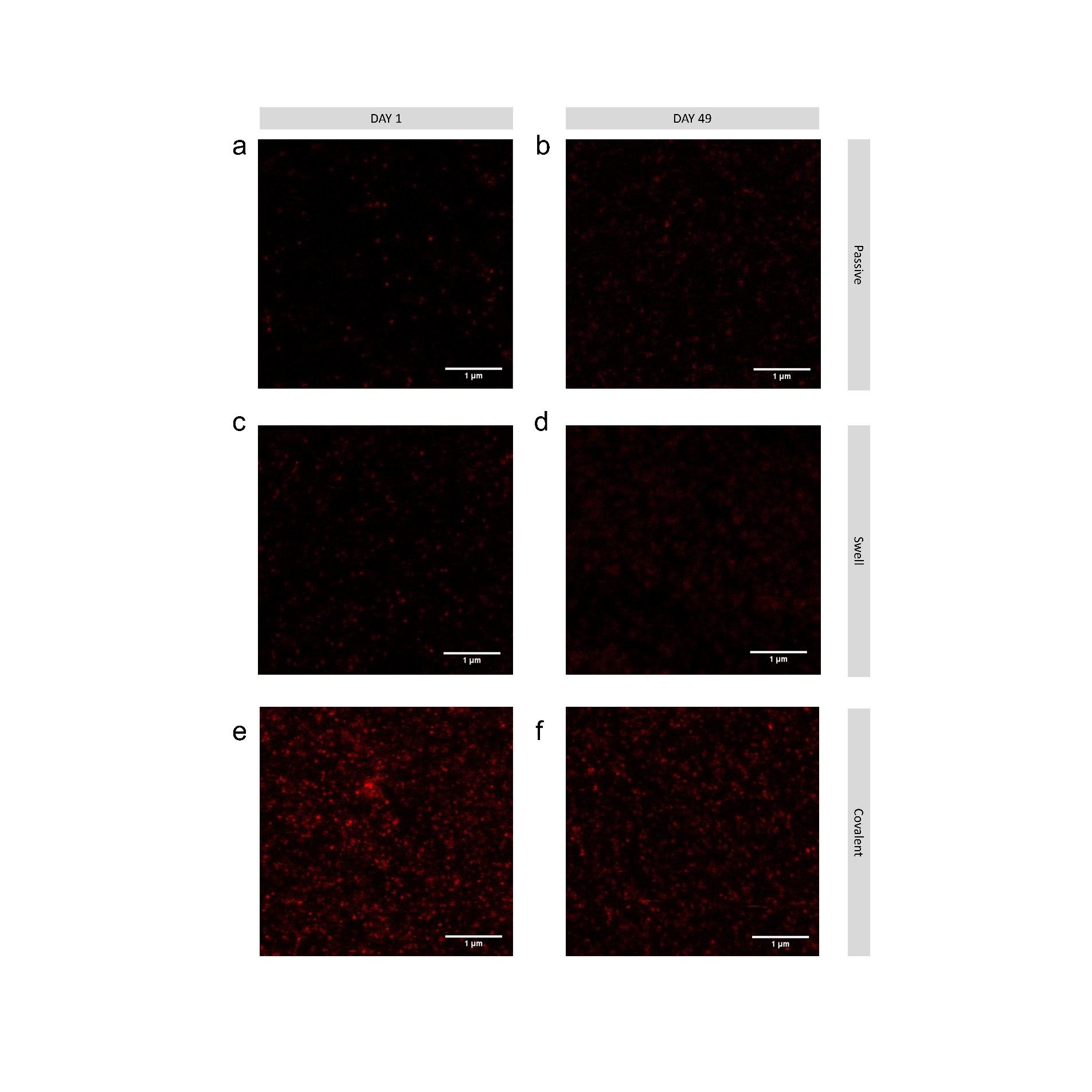

**Supplementary Figure 5:** Comparison between images of 100 nm polystyrene beads initially and after 49 days in water for passive (a,b), swell (c,d), and covalent (e,f) labeling with Atto 647N.

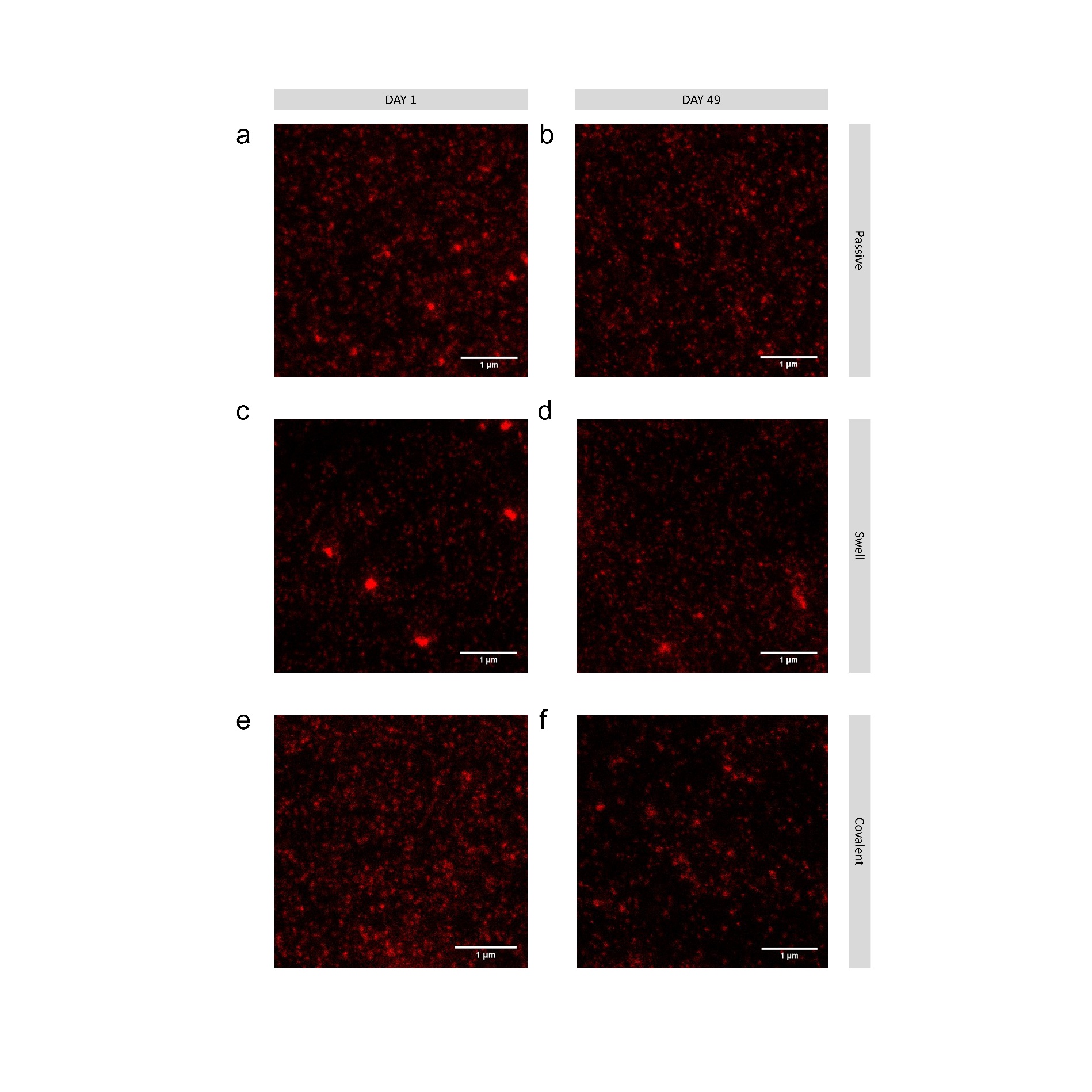

**Supplementary Figure 6:** Comparison between images of 100 nm polystyrene beads initially and after 49 days in oil for passive (a,b), swell (c,d), and covalent (e,f) labeling with Atto 647N.

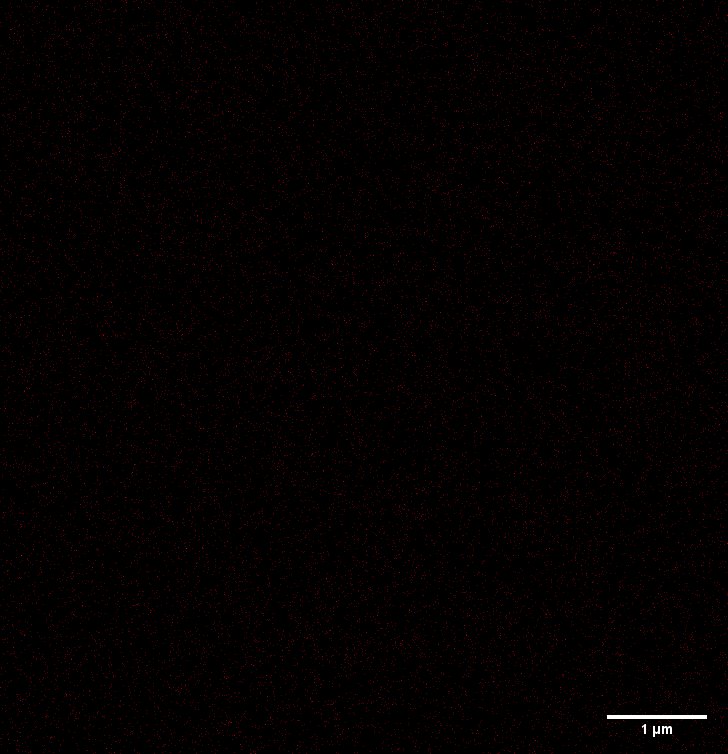

**Supplementary Figure 7:** Free dye dialyzed control STED image. No detectable signal from dialyzed solution, suggesting that free dye was largely removed with the dialysis step.

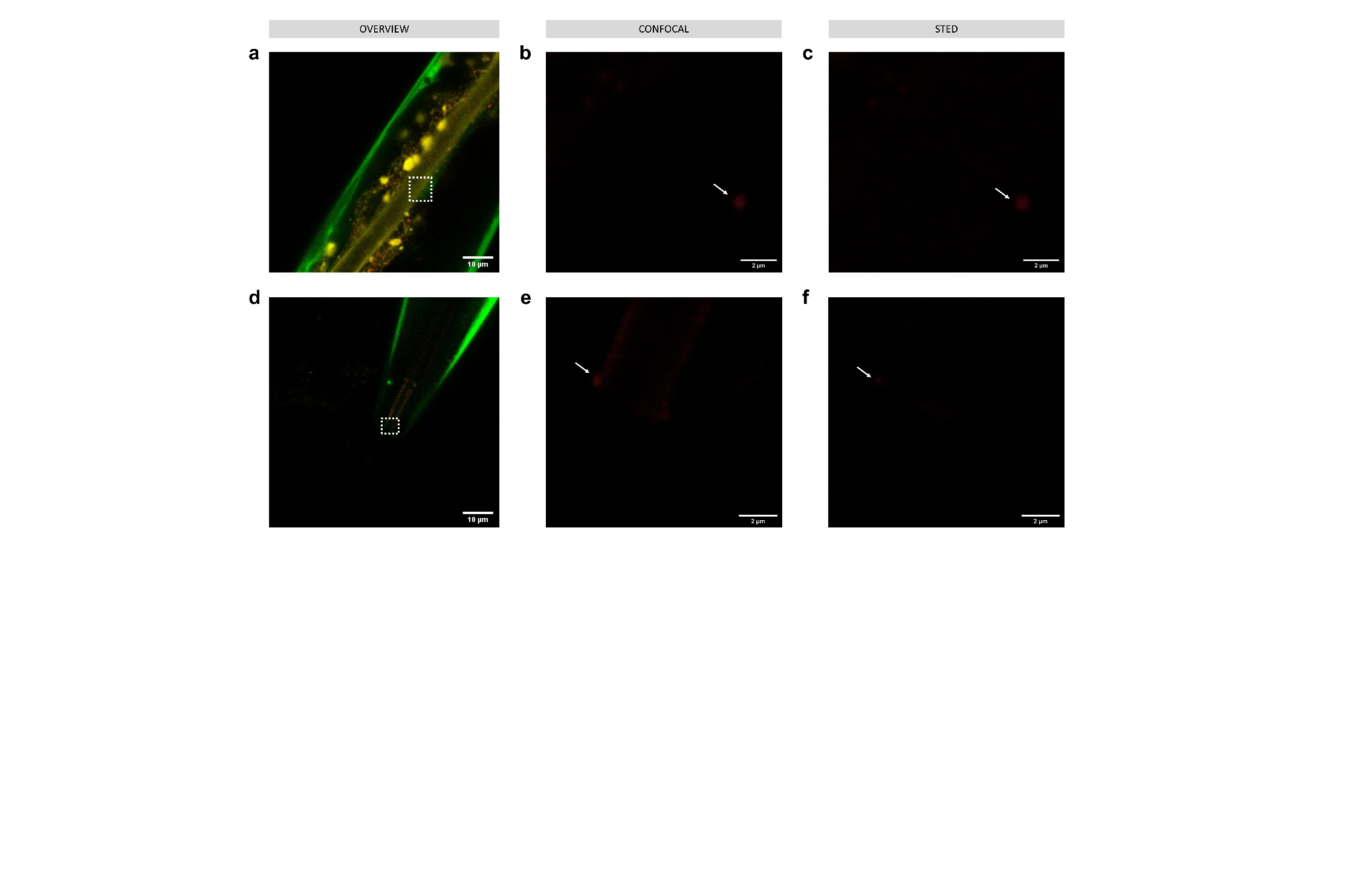

**Supplementary Figure 8:** Imaging *C. elegans* KWN117 adult exposed to dialyzed dye solution (no nanoparticles) expressing GFP (green) in the body wall and mCherry (yellow) in the apical intestinal membrane*.* Confocal overview, and high resolution of confocal and STED images of scanned area indicated by white boxes for parts of the digestive track in the intestine (a,b,c) and mouth (d,e,f). Note that while nanoparticles are not seen there is still some signal, likely from autofluorescence, that would have potentially been misinterpreted as labeled nanoplastic particles, due to a truer image size, but for the resolution provided by STED microscopy. Arrows indicate potential false positive signal from confocal imaging where STED imaging reveals that the signal is inconsistent with 50 nm particles.

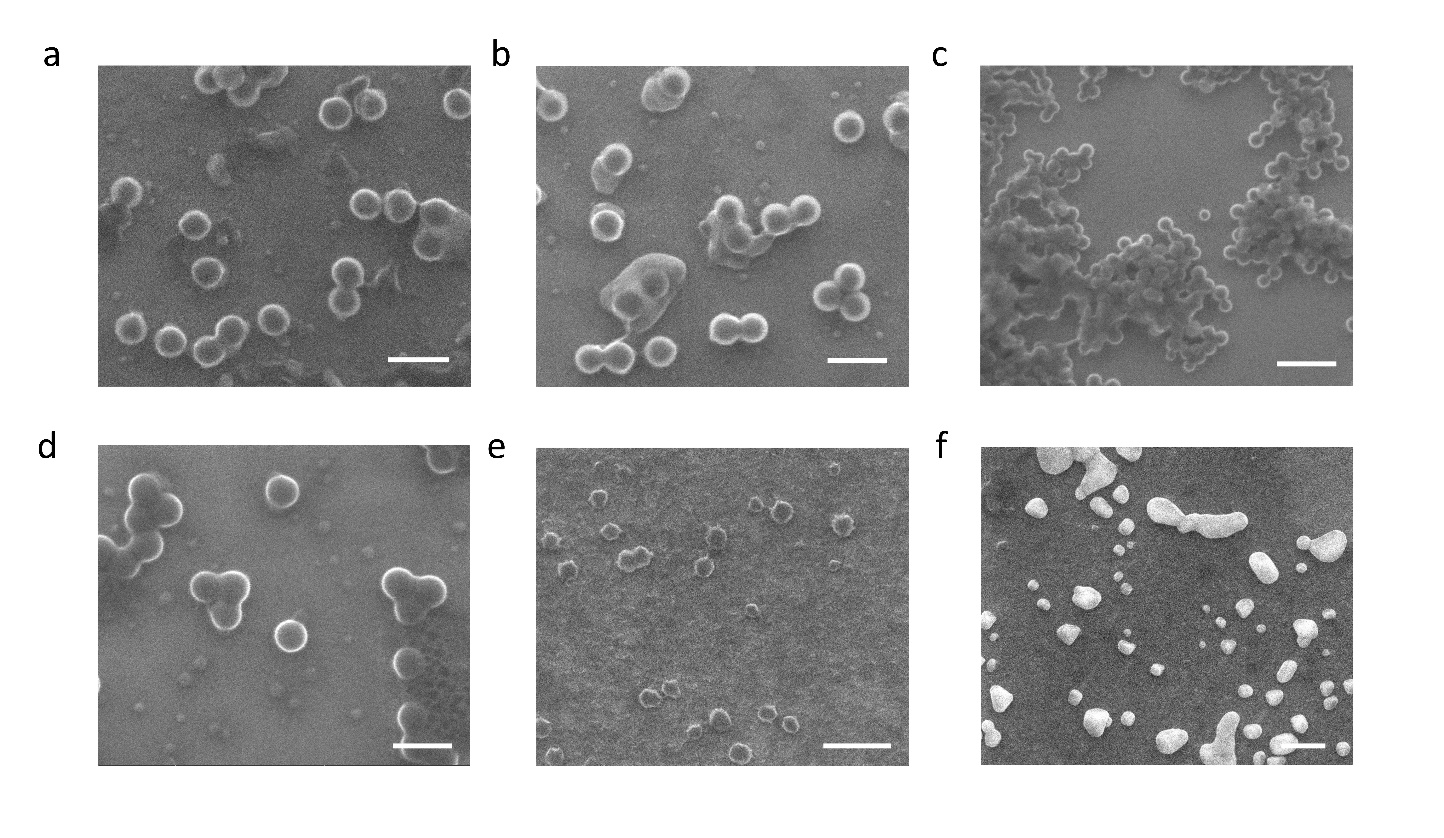

**Supplementary Figure 9:** SEM images of samples including (a) 100 nm plain polystyrene beads – scale bar is 200 nm, (b) 100 nm amine-modified polystyrene beads – scale bar is 200 nm, (c) 50 nm plain polystyrene beads – scale bar is 200 nm, (d) PTFE particle suspension – scale bar is 200 nm, (e) debris from a heated polystyrene plate – scale bar is 400 nm, and (f) debris from sanding a polystyrene Petri dish – scale bar is 500 nm.
